# Analysis of Gene Expression Profiles to study Malaria Vaccine Dose Efficacy & Immune Response Modulation

**DOI:** 10.1101/2021.08.05.454986

**Authors:** Supantha Dey, Harpreet Kaur, Mohit Mazumder, Elia Brodsky

## Abstract

Malaria is a life-threatening disease, and the Africa is still one of the most affected endemic regions despite years of policy to limit infection and transmission rates. Further, studies into the variable efficacy of the vaccine are needed to provide a better understanding of protective immunity. Thus, the current study is designed to delineate the effect of the different vaccination doses on the transcriptional profiles of subjects to determine its efficacy and understand the molecular mechanisms underlying the protection this vaccine provides. Here, we used gene expression profiles of pre and post-vaccination patients after various doses of RTS,S based on 275 and 583 samples collected from the GEO datasets. At first, exploratory data analysis based Principal component analysis (PCA) shown the distinct pattern of different doses. Subsequently, differential gene expression analysis using edgeR revealed the significantly (FDR <0.005) 158 down-regulated and 61 upregulated genes between control vs. Controlled Human Malaria Infection (CHMI) samples. Further, enrichment analysis of significant genes using Annotation and GAGE tools delineate the involvement of *CCL8*, *CXCL10*, *CXCL11*, *XCR1*, *CSF3*, *IFNB1*, *IFNE*, *IL12B*, *IL22*, *IL6*, *IL27*, etc.,genes which found to be upregulated after earlier doses but downregulated after the 3^rd^ dose in cytokine-chemokine pathways. Notably, we identified 13 cytokine genes whose expression significantly varied during three doses. Eventually, these findings give insight to the dual role of cytokine responses in malaria pathogenesis and variations in their expression patterns after various doses of vaccination involved in protection.

## Introduction

Malaria remains a well-known and life-threatening disease in many tropical and subtropical countries. Currently, there are over 100 countries and territories where the risk of malaria transmission is present [6]. These countries are visited by more than 125 million international travelers every year [1]. Moreover, Africa has faced 94% of all malaria cases in 2019. There were almost 229 million estimated malaria cases worldwide in the same year. And the number of deaths from malaria stood at more than 400,000. Malaria is known to be caused by *Plasmodium* parasites. These parasites can infect female Anopheles mosquitoes and spread to people through the biting from these mosquitoes. Among the five parasite species that cause malaria in humans, two species- *P. falciparum* and *P. vivax* bear the highest threat. *P. falciparum* also accounted for almost 99.7% of malaria cases in the African region and nearly half of WHO South-East Asia Region cases. [2]

Most malaria deaths occur in children, and they are dominated by three syndromes: severe anemia, cerebral malaria, and respiratory distress. These syndromes can occur separately, or in combination: [3] One of the elementary features of *P. falciparum* is the induction of host inflammatory responses that contribute to disease severity and are associated with lethal outcomes. [4] Specifically, systemic levels of some pro-inflammatory cytokines are correlated with severity and death from malaria. [5]

Even though the disease appears in documented reports as early as 2700 B.C., malaria vaccine development entered a new milestone only in 2015. [6]. The European Medicines Agency positively reviewed the pre-erythrocytic Plasmodium falciparum candidate RTS,S vaccine and marked the first human anti-parasite vaccine to pass the regulatory examination. [7] This vaccine provided protection against infection in controlled human malaria infection (CHMI) studies [22–24]. It can also prevent life-threatening malaria and reduce the need for transfusion of blood. [8]

Although malaria has been studied in detail, insufficient attention has been paid to how malaria vaccination is associated with several gene expression changes,contributing to increased protection. The efficacy of the malaria vaccine is still not at a desirable level and needs improvement if we want to eradicate malaria. [9] In the current study, we focused on how the various doses of malaria RTS,S/AS01 vaccine facilitates protection and gene expression changes. Though we discussed multiple doses, we were primarily focused on a dataset from the 3rd dose. It could give us a better overview of gene expression changes as delayed doses are found to increase the chance of protection against malaria. [10] RTS,S/AS01 vaccination has been observed to be significantly associated with upregulation and downregulation in several gene expressions. [11]. Our study would help us conclude how several gene upregulation & downregulation after the vaccination is different from dose to dose and how cytokine’s dual role in protection & pathogenicity in malaria is crucial to investigate. Moreover, it will investigate how the absence of negative feedback control in pathophysiologic situations is responsible for impairing cytokine network homeostasis and contribute to local pathogenesis. [12].

## Methods

Figure 1 provides us an overall idea of the workflow of the research and methods we used to obtain the results.

**Figure 1:**
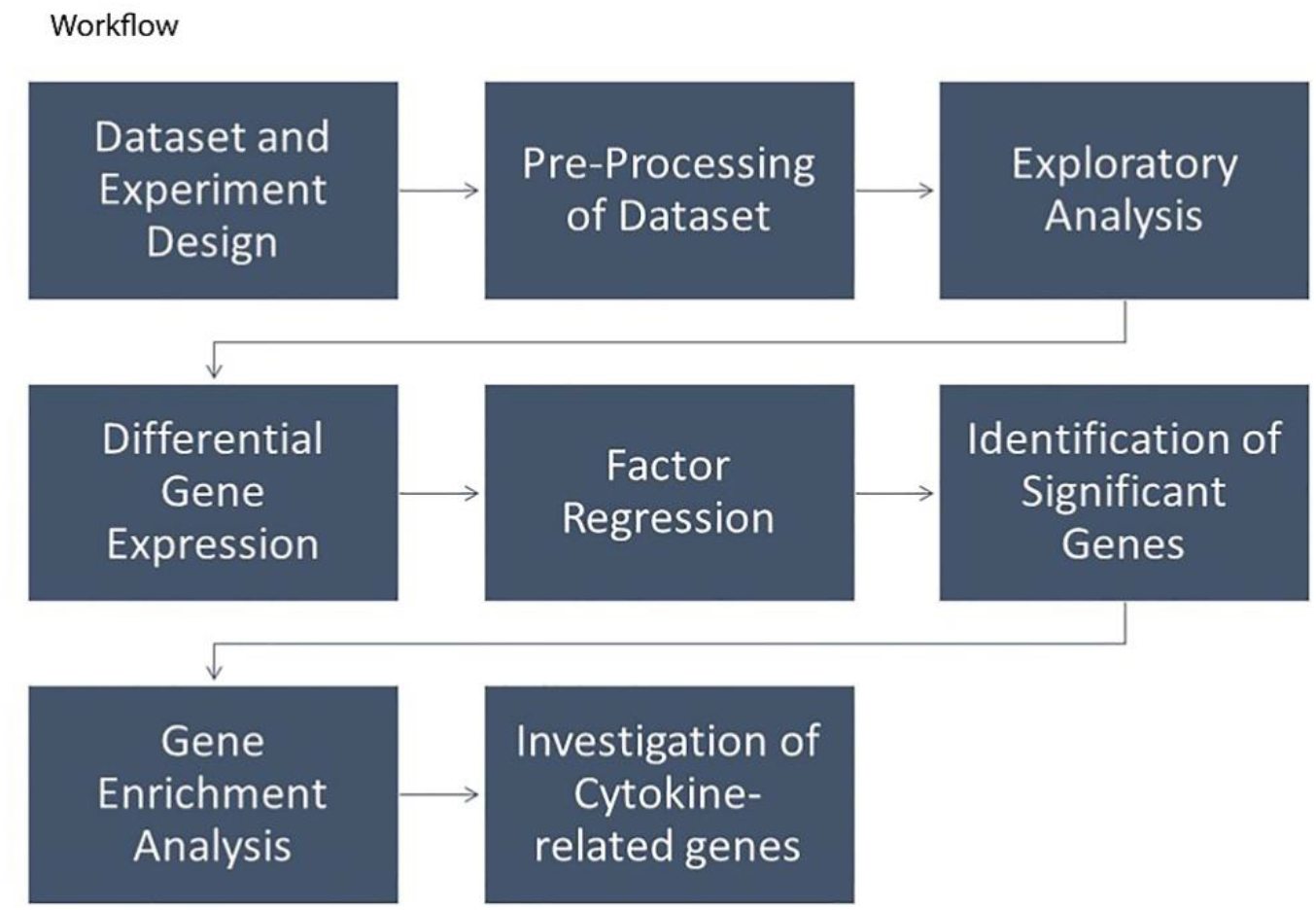
Workflow of the Research Process.

### Dataset & Experimental Design

Two datasets were obtained from GEO with accession numbers GSE102288 and GSE89292. These datasets were published as a BioProject on the NCBI with the accession number PRJNA397222 and PRJNA351258. The total number of samples was 275 in GSE102288 and 583 in GSE89292. [13] [14]. An overview of the datasets is given in Table 1.

**Table 1:** Original Dataset source, design, analysis method and sample number.

| Dataset | Organism | Experiment Type | Sample Source | Analysis Method | Sample Number. |
| --- | --- | --- | --- | --- | --- |
| GSE102288 | Homo Sapiens | Expression profiling by high throughput sequencing | PBMC samples | RNAseq analysis. | 275 |
| GSE89292 | Homo Sapiens | Expression profiling by array | PBMC samples | RNAseq analysis. | 583 |

### Datasets for the Identification of Differentially Expressed Genes

We partitioned datasets for two comparative analyses: The GSE102288 dataset was analyzed to compare control vs. CHMI samples, and GSE89292 was analyzed to identify gene expression patterns after the first two doses. Illumina HiSeq 2000 was the used platform for GSE102288,and Affymetrix Human Genome U133 Plus 2.0 Array was used in GSE89292.All datasets were curated so that only human tissue samples remain in the dataset. Furthermore, Probe ID mapped to gene symbols in the GSE89292 dataset was extracted from the respective platform file. Finally, dataset matrices were prepared for various analyses.

### Pre-Processing of Datasets

The GSE102288 dataset contains the FPKM value for 15, 680 genes. On the other hand, GSE89292 is a microarray data containing RMA normalized value. In the case of the Illumina dataset (GSE102288), FPKM values are converted into log2 values using the T-bioinfo server pipeline (Figure 2). In the case of Affymetrix datasets (GSE 89292), the average of multipleprobes was computed that correspond to a single gene using the average function in Excel.Ensembl transcript I.D.s were mapped to the gene symbols using the T-bioinfo Server’s annotation pipeline.

**Figure2:**
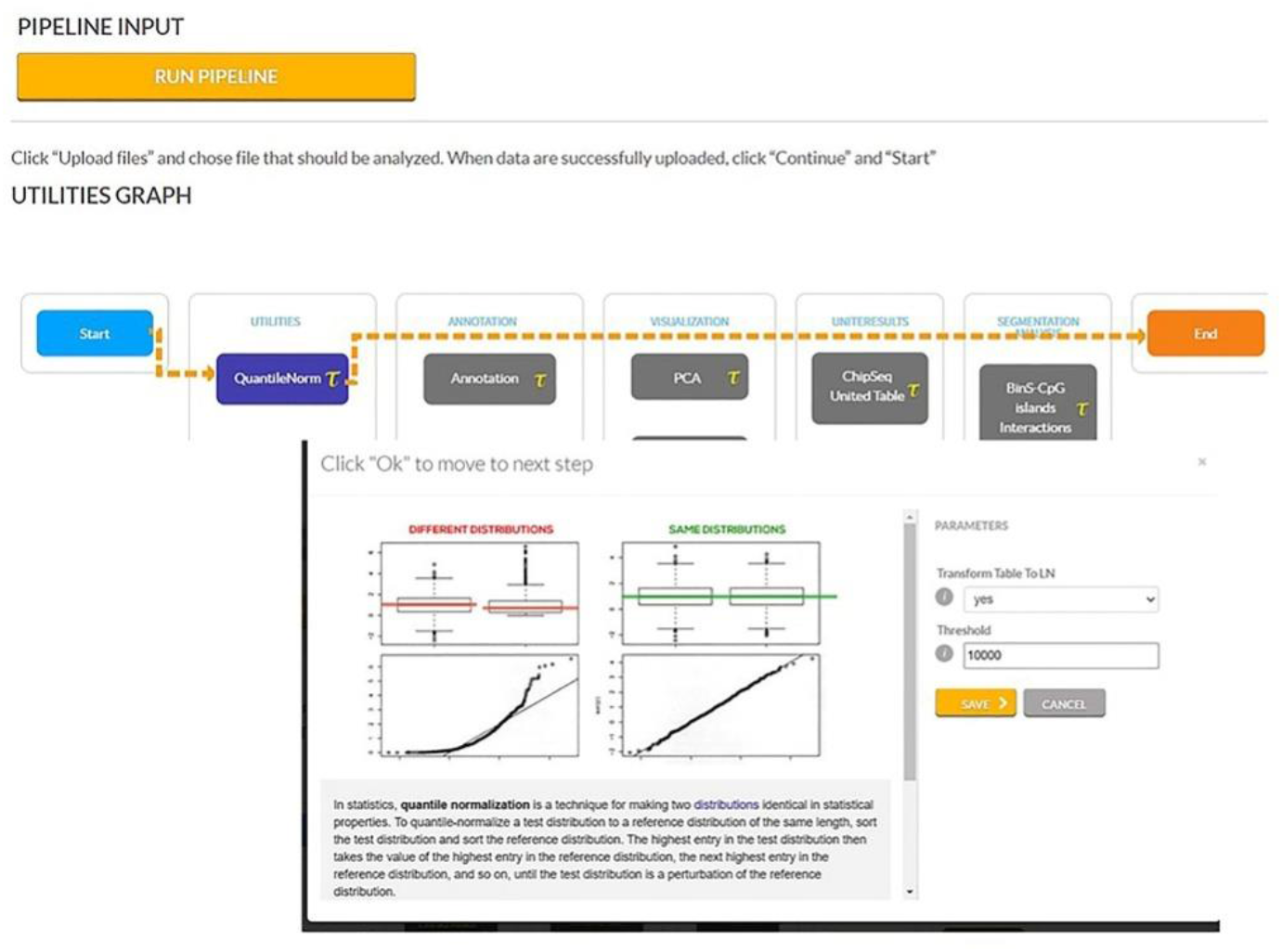
Quantile Normalization Pipeline.

### Exploratory Analysis

In order to explore the patterns of the data, principal component analysis (PCA) was performed using the T-bioinfo Server (https://server.t-bio.info/) on all the 275 samples based on the gene expression profiles of the samples. Besides, PCA and Hierarchical Clustering (distance: Euclidean, linkage: ward.D2) were also performed with only significant genes to assess their discriminant potential [15] using the T-bioinfo Server.

### Differential Gene Expression Analysis

We performed differential gene expression analysis using the EdgeR [16] tool integrated on the T-Bioinfo Server to select genes that were significantly differentially expressed in Pre-vaccination (control) vs. Day of Challenge (CHMI) samples (Figure 3a). The T-Bioinfo Server was used for this purpose. Furthermore, p-value and Log2 fold change values obtained from edgeR results were used to generate a volcano plot (Figure 4) for control vs. CHMI samples. Eventually, a heatmap was generated for the top 50 protein-coding upregulated and downregulated genes. [17]. Only those genes were considered as significant genes with the False Discovery Rate (FDR) < 0.005 and log2 fold change at > ±1. We applied this threshold for both datasets.

**Figure 3:**
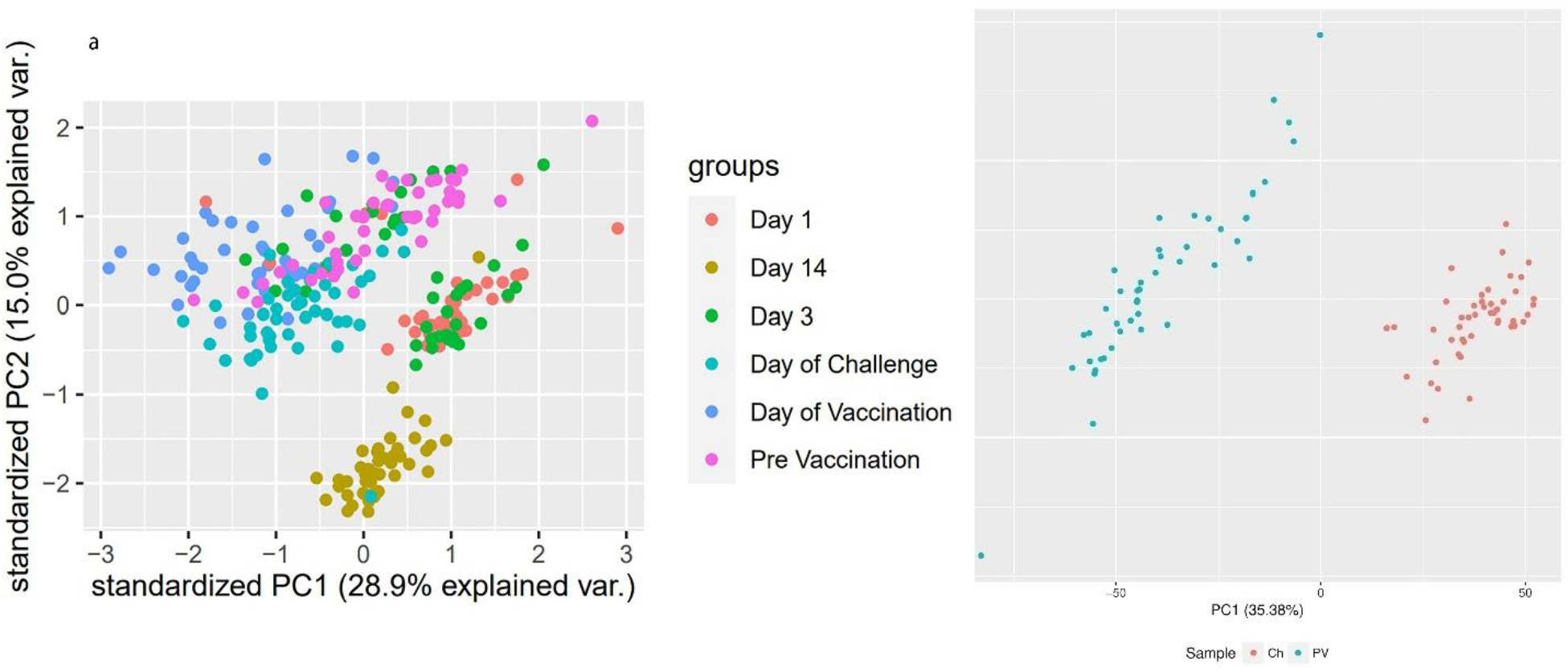
(a) PCA plot for all sample groups in the complete dataset and (b) PCA plot for control vs. CHMI samples.

**Figure 4:**
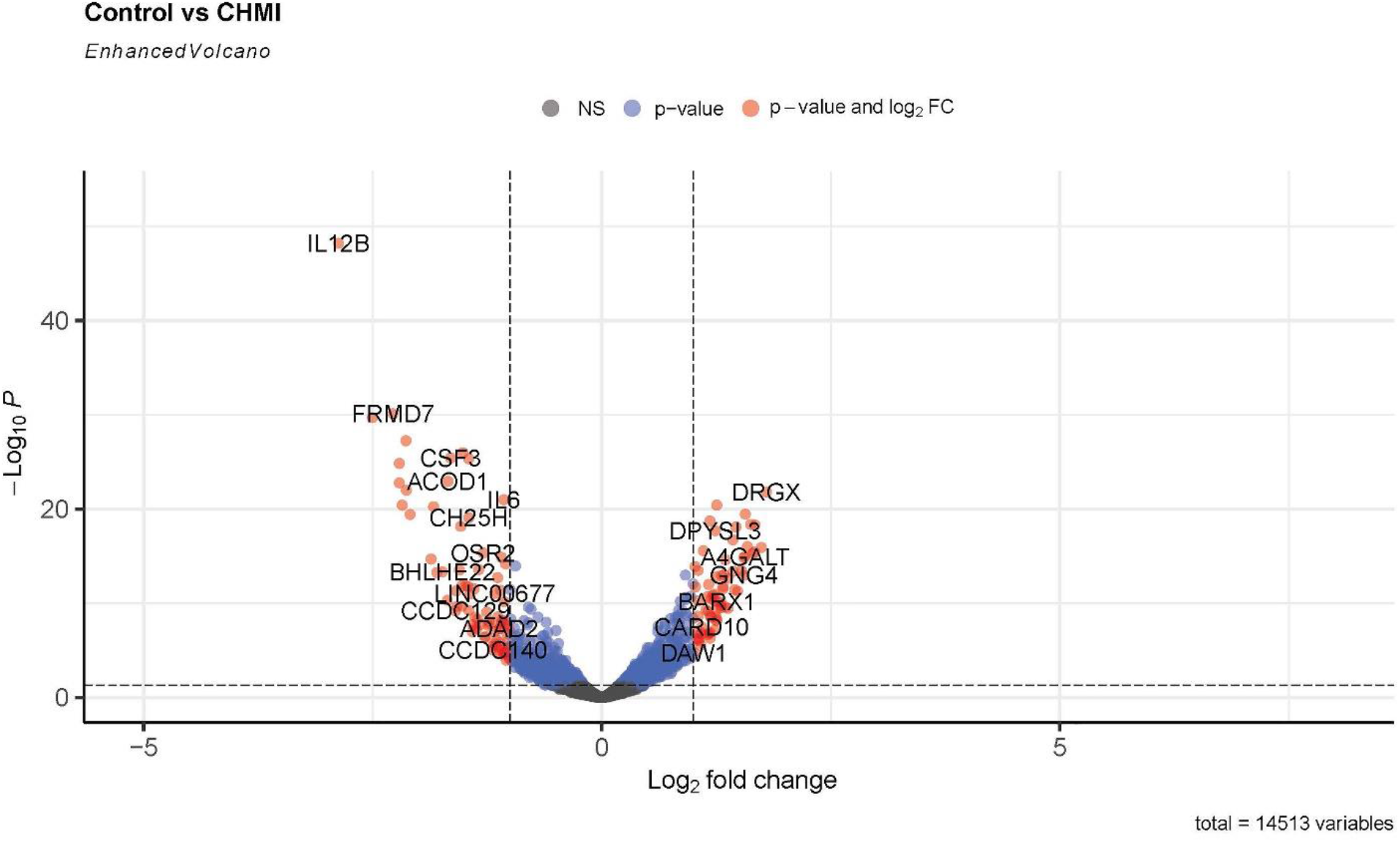
Enhanced volcano plot for control and CHMI samples for all annotated genes in R.

### Factor Regression analysis

Next, we conducted a factor analysis to interpret the relationship between multiple variables and the malaria vaccine efficacy. (Supplementary Figure 2) Correlation between gender status, protection status, and gene expression was investigated. Our study also helped us to find out how other factors are involved in protection status and which genes are significantly expressed in protected samples. [50]

### Gene Enrichment Analysis

To understand how the significant genes are related to protection, we assessed the biological and molecular functions of the significant genes with the help of annotation. We used Database for Annotation, Visualization, and Integrated Discovery (DAVID) for annotation. [18] Additionally, we used the Kyoto Encyclopedia of Genes and Genomes (KEGG) and Bioplanet for pathway analysis. [19] [20] Literature search was done for the top significant genes, and cytokine or inflammation-associated genes provided us a better understanding.

## Results

### Detection and visualization of variation in Data

The exploratory data analysis based on the PCA plot revealed the separate clusters for various sample groups. Principal component 1 (PC1) represents 28.9% variance and PC2 represents 15.0% variance of the data (Figure 3a). The figure illustrates how various samples fall in different clusters and overlaps as well. Interestingly, here, we observed Day 14 samples forming a distinct cluster. Day 1 and day 3 post 3rd vaccination samples remain together in a cluster as they indicate the early expression of genes after the dose. The CHMI (day of the challenge) and control (pre-vaccination) samples also form distinct clusters. So, it is most likely that the gene expression pattern after malaria vaccine doses can indicate how the patient responds to the vaccine. Furthermore, PC1 represents 35.38% variance, and PC2 represents 13.73% variance of the data in control vs. CHMI samples only and falls into different clusters. (Figure 3b).

### Identification of Differentially Expressed Significant Gene

Since the PCA plot showed clear, distinct clusters for samples of control and CHMI, next, we identified 219 significantly differentially expressed genes between control and CHMI samples based on EdgeR. Among them, 158 genes were found to be down-regulated, and 61 genes were found to be upregulated in control vs. CHMI samples. Further, we generated a volcano plot that included all the genes (Figure 4). The volcano plot represents significant genes between control and CHMI groups. Furthermore, to identify a manageable subset of genes, we selected only the top 50 differentially expressed genes, including the top 25 upregulated and top 25 downregulated in control vs. CHMI samples (Table 2). Notably, here we have selected only protein-coding and excluded the non-protein-coding genes. Although, noncoding RNA genes have been found to help evade human immune attack through switching in expression between variants of var family genes and increase the severity of malaria infection. but not in inflammation and protection. [40] [39] Since our study primarily focused on the human inflammatory gene expression and its association with protection or elevated severity of malaria infection; therefore we excluded the noncoding genes.

**Table 2:** Top 50 significantly expressed protein-coding genes (P-value<0.05, FDR <0.005, and log2 fold change > ±1).

| Downregulated Genes |  |  | Upregulated Genes |  |  |
| --- | --- | --- | --- | --- | --- |
| Gene Symbol | logFC | FDR | Gene Symbol | logFC | FDR |
| IL12B | -2.87295 | 9.85E-45 | DRGX | 1.798081 | 1.35E-19 |
| FRMD7 | -2.27791 | 4.06E-27 | CHST3 | 1.743642 | 4.84E-14 |
| IFNB1 | -2.20904 | 2.05E-22 | KIAA1644 | 1.674141 | 2.76E-16 |
| IL36G | -2.1776 | 2.89E-18 | CACNA1G | 1.669566 | 1.44E-13 |
| CXCL11 | -2.13596 | 1.78E-24 | TNFAIP8L3 | 1.65029 | 1.80E-13 |
| CCL8 | -2.13175 | 9.23E-20 | NECTIN4 | 1.627141 | 2.36E-16 |
| RANBP3L | -1.86129 | 6.74E-13 | RHBDL3 | 1.590967 | 4.04E-14 |
| IL22 | -1.80012 | 1.40E-11 | SMAD6 | 1.567576 | 2.30E-17 |
| BHLHE22 | -1.73922 | 1.23E-11 | A4GALT | 1.565569 | 4.01E-13 |
| ACOD1 | -1.67863 | 1.28E-20 | GNG4 | 1.553002 | 2.61E-11 |
| CSF3 | -1.64862 | 6.61E-23 | HR | 1.550488 | 4.74E-13 |
| CCDC129 | -1.59496 | 7.71E-08 | MAFA | 1.526466 | 9.91E-12 |
| UNC5C | -1.589 | 8.93E-10 | LMX1B | 1.483051 | 9.46E-10 |
| IFNE | -1.58164 | 3.03E-08 | SEMA3F | 1.466303 | 3.83E-16 |
| C1QTNF1 | -1.55235 | 8.45E-12 | CBARP | 1.452397 | 6.85E-10 |
| PLA1A | -1.539 | 3.60E-16 | HOXB6 | 1.435488 | 1.08E-11 |
| GXYLT2 | -1.52936 | 3.14E-10 | RASAL1 | 1.431945 | 8.37E-15 |
| HSD11B1 | -1.52447 | 2.82E-08 | UBE2QL1 | 1.383004 | 4.92E-11 |
| CXCL10 | -1.51384 | 2.10E-23 | CA12 | 1.378161 | 4.43E-08 |
| RHCG | -1.48479 | 2.99E-10 | NCMAP | 1.351544 | 3.31E-08 |
| KCNJ2 | -1.45138 | 7.38E-23 | FAM124A | 1.34904 | 7.86E-13 |
| SLC6A7 | -1.45109 | 6.99E-08 | MMP2 | 1.333238 | 2.41E-11 |
| CH25H | -1.45018 | 4.82E-17 | GAD1 | 1.325456 | 3.36E-10 |
| OR52W1 | -1.40885 | 1.71E-06 | TUBB3 | 1.324794 | 5.41E-10 |
| IL27 | -1.39053 | 5.41E-10 | RGMA | 1.307805 | 2.88E-11 |

### PCA plot based on only significant genes

Next, the PCA plot based on only significant genes between control and CHMI shows a clear distinction among the samples of these groups. Principal component 1 (PC1) represents 78.74% variance, and PC2 represents 3.64% variance of the data. (Supplementary Figure 3).

### Hierarchical Clustering and Heatmap for Top 50 Genes Show Clear Distinctive Pattern

Next, to assess the capability of the top 50 significant genes in distinguishing control & CHMI samples, we performed H-Clustering with this set of genes. The dendrogram represents the two clear, distinct clusters of control and CHMI samples (figure 6b). Heatmap represents the expression pattern of the top 50 top genes between control vs. CHMI samples (Figure 6a), illustrating variation in expression. Among the significantly expressed genes after the 1^st^ and 2^nd^ dose of vaccination, we identified the same cytokine & inflammation-related genes that were also significantly expressed after the 3^rd^ dose. In order to derive their correlation and visualize the expression pattern, we plotted a line chart (Figure 5a and 5b). After giving the first dose, there is upregulation of the inflammatory genes and both up and down-regulation after the second dose (Figure 5a). Interestingly, significant downregulation occurs to those same genes after applying the third vaccination dose (Figure 5b). The gene expression pattern changed through various doses of vaccination.

**Figure 5:**
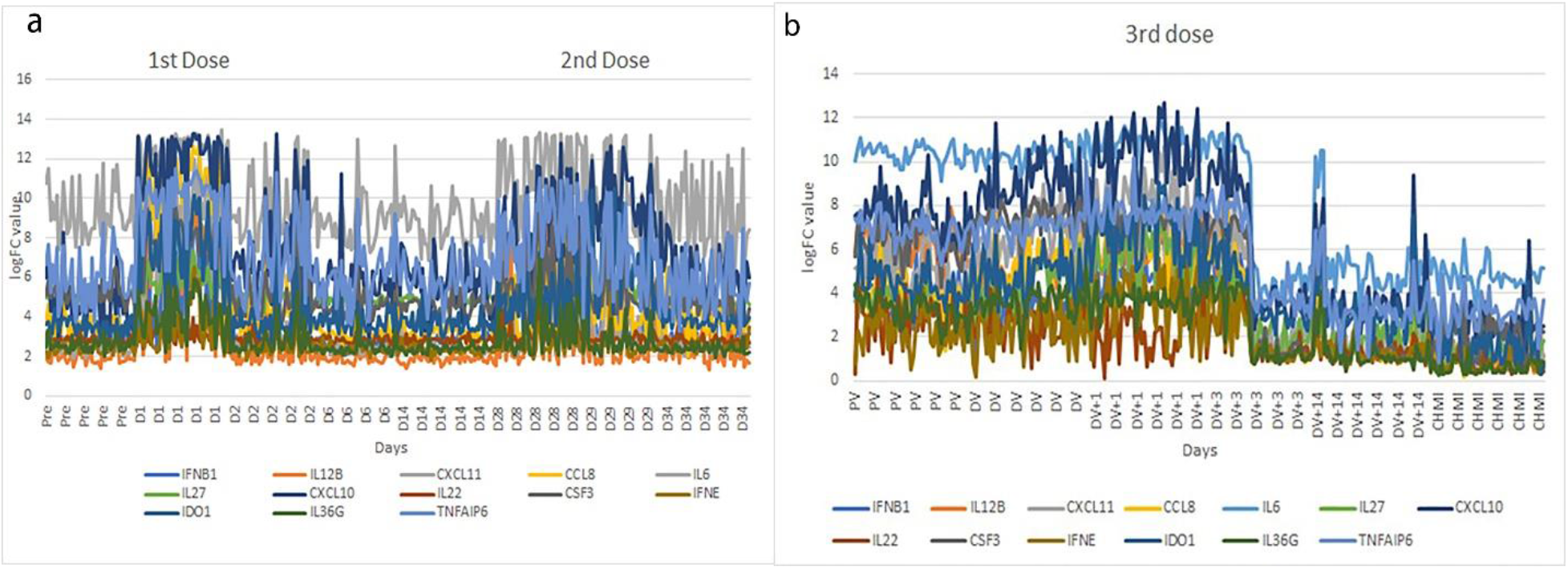
Inflammatory or cytokine gene expression pattern in (a) 1st and 2nd dose and (b) 3rd dose distinct up or down-regulation.

**Figure 6:**
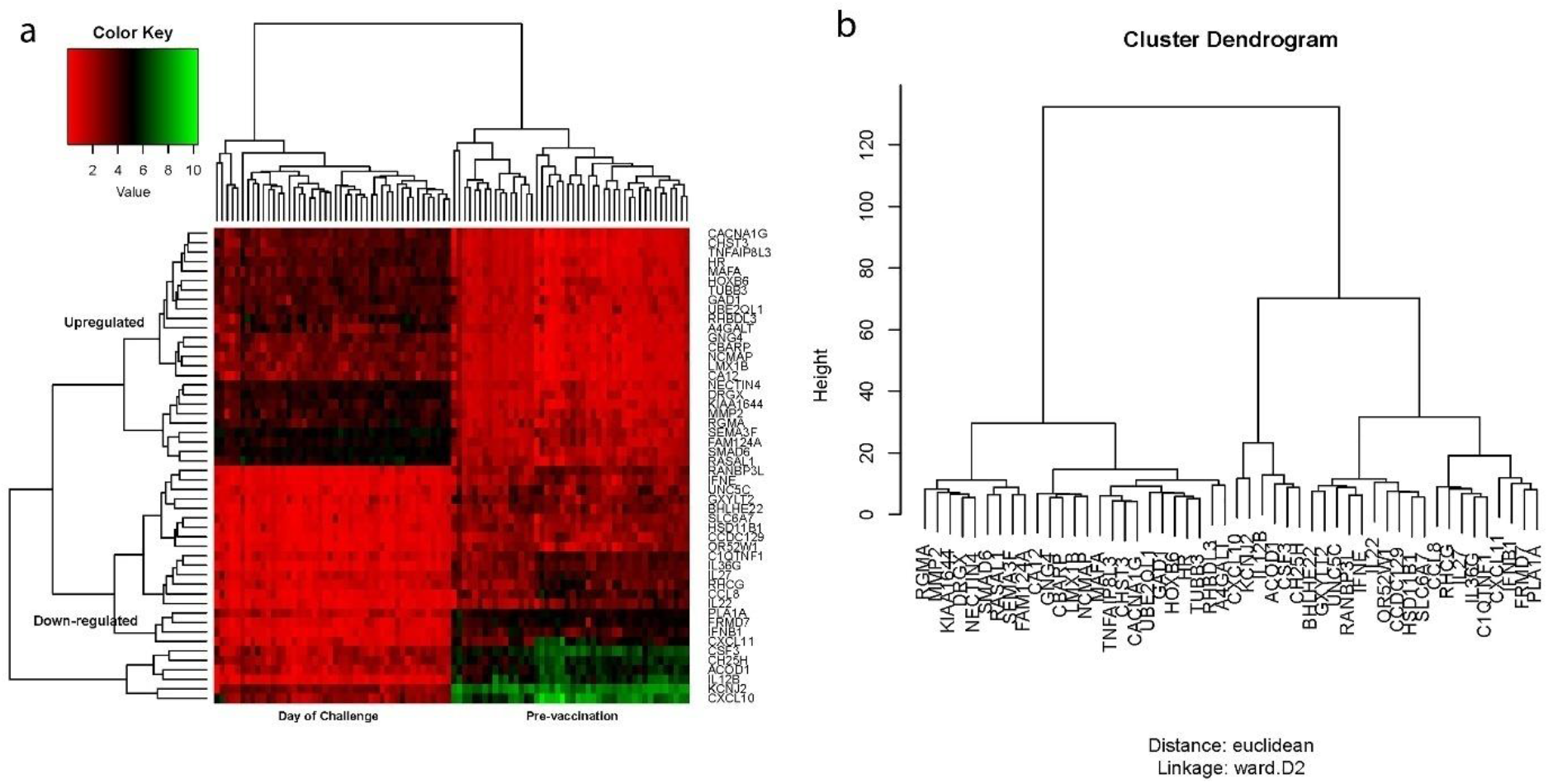
(a) Heatmap & (b) Dendrogram generated by Hierarchical clustering based on top 50 significantly differentially expressed genes, illustrating the distinct clusters of control and CHMI day samples.

### Expression of Inflammatory Cytokine Genes in All Doses

Cytokines are the key players of the immune system. Next, to understand the expression pattern of the cytokines during different doses, we investigated the expression pattern for significantly expressed genes with cytokine G.O.function in control vs. CHMI samples. We identified 13 significantly differentially expressed cytokine genes that were expressed after all three doses. These cytokine genes showed a distinctive expression pattern after different doses. (Figure 5a and 5b)

### Correlational with Other Factors of Interests

Next, we wanted to see whether the gender of the samples is a significant factor in gene expression regulation. Here, we identified 10 protein-coding genes that were indeed significantly associated with the gender (males and females) on control vs. CHMI comparison (Supplementary Table1). A PCA plot was also generated to visualize the differences (Supplementary Figure 4). We further looked for evidence if gender was correlated with protection. No significant genes were found to provide any evidence of gender to be related to protection. Subsequently, we identified13 coding genes significantly differentially expressed in protected vs. non-protected samples on the 3^rd^ vaccination day vs. CHMI samples. (Supplementary Table 2) The exploratory data analysis failed to show enough differences between samples while comparing 3^rd^ vaccination vs. CHMI samples, but further analysis is necessary for identifying the importance of the significantly expressed genes after 3^rd^ dose on protected vs. non-protected samples.

### Gene Set Enrichment Analysis

As malaria severity is often associated with the over-expression of inflammatory genes, their expression patterns and changes were a major point of interest in our study as we wanted to find out how vaccination doses affect them.

Gene set enrichment analysis was performed for 158 upregulated, 61 downregulated significant genes in CHMI vs. control samples of the 3rd dose on DAVID. Also, functional annotation and clustering for these significantly expressed genes were performed. Top hits with the downregulated genes indicate the enrichment in gene ontology terms associated with inflammatory response, cellular response to lipopolysaccharide, cytokine, chemokine mediated signaling pathway, cell-cell signaling, positive regulation of leukocyte chemotaxis, etc. Functional clustering showed 9 clusters with 83 DAVID IDs. The top three clusters were all involved in cytokine activity and had enrichment scores of 4.42, 3.19, and 3.15, respectively. These enrichment scores are measured by the geometric mean of the EASE Scores (modified Fisher Exact) [68]. Here, a higher score for a group is an indication of their more critical (enriched) roles. [48]. Similarly, gene enrichment analysis with the upregulated genes for. CHMI vs. controlshowed the gene ontology enriched terms were associated with intramembranous ossification, negative-regulation of calcium ion-dependent exocytosis, epidermis development, angiogenesis, chemical synaptic transmission, positive regulation of calcium ion-dependent exocytosis, cell-cell signaling, etc. Also, functional clustering indicated 9 clusters with 68 DAVID IDs, where top ones were associated with glycoprotein, glycosylation site: N linked, and pathways in cancer. Enrichment scores were 1.88, 1.62 & 1.39 for the top three clusters. We also analyzed a dataset from the 1^st^ dose, which showed association with inflammatory response, cytokine, and chemotaxis. Interestingly, CCL7, CXCL1, CXCL11, etc. cytokine genes were upregulated in this case whether they were downregulated after the 3^rd^ dose.

Besides, gene enrichment analysis of significantly differentially expressed genes in protected vs. non-protected showed the enrichment in gene ontology terms,including cell-matrix adhesion and association of signal and glycoprotein. [21]

### KEGG Pathway Analysis

KEGG Pathway analysis revealed genes, such as *CCL8*, *CXCL10*, *CXCL11*, *XCR1*, *CSF3*, *IFNB1*, *IFNE*, *IL12B*, *IL22*, *IL6*, *IL36G*, *IL27*, etc. were involved in cytokine-cytokine receptor interaction (Figure 7). Moreover, *CSFE*, *IFNB1*, *IFNE*, *IL12B*, *IL22*, *IL6* genes were associated with the JAK-STAT signaling pathway. (Supplementary figure 5)

**Figure 7:**
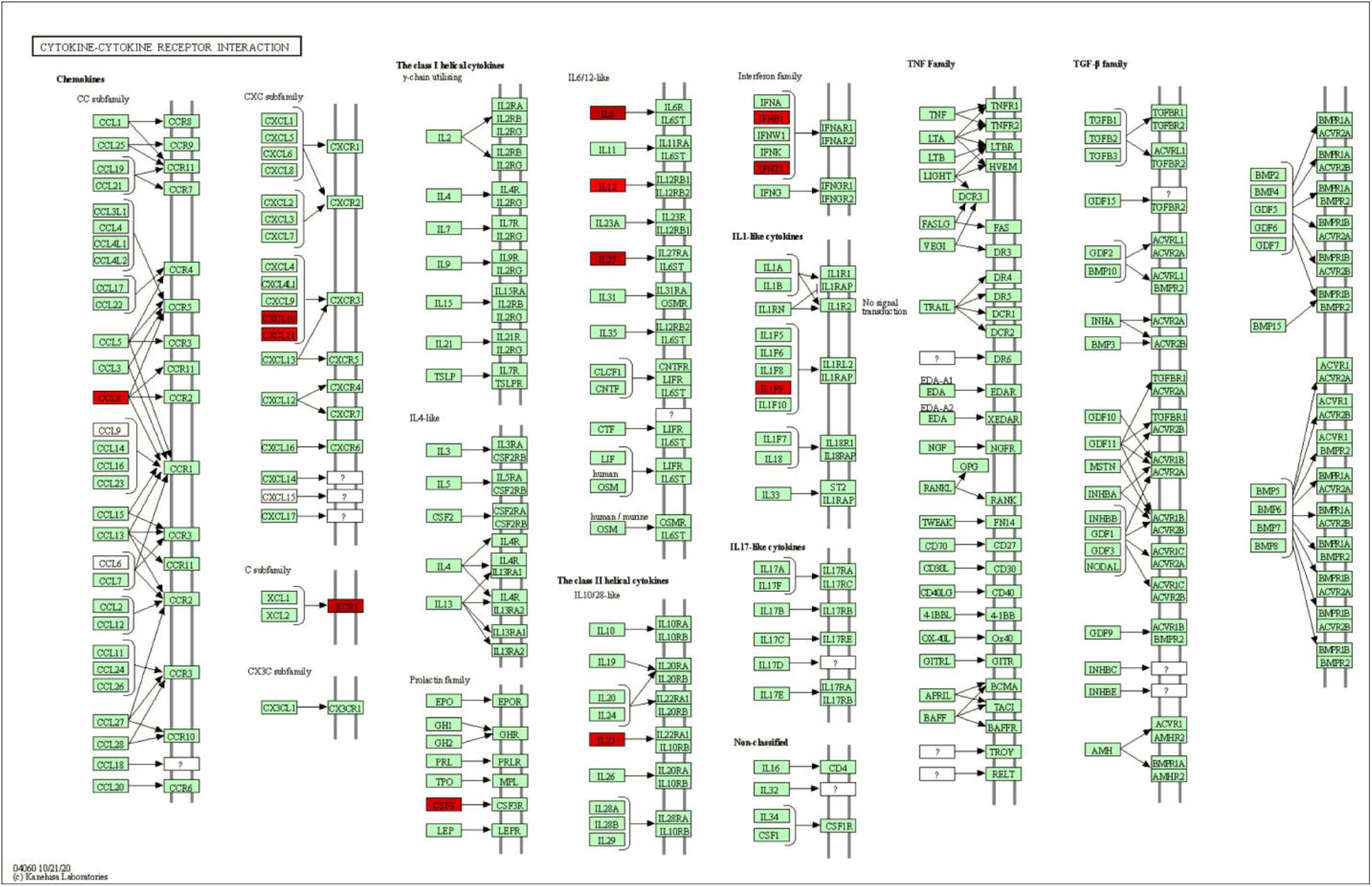
KEGG Cytokine-Cytokine Receptor Interaction Pathway involving significant genes in control and CHMI samples.

On the other hand, the upregulated genes, such as *GNG12*, *GNG4*, *WNT9A*, *EDNRB*, *FGF18*, *LAMA3*, *MMP2*, etc., were found in pathways in cancer (supplementary figure 6).

## Discussion

There are two main determinants of severe malaria: sequestration of parasitized RBC and surge of pro-inflammatory responses. [37] Imbalanced pro and anti-inflammatory immune responses have been found to trigger immune-induced pathology and remain one of the leading causes of cerebral malaria pathogenesis, which might be further amplified by sequestration. [27]. Moreover, systemic cytokine levels are correlated with disease severity in malaria as well as sepsis. [26] Thus, it is necessary to examine how multiple vaccination doses change the pattern of inflammatory responses and induce protection upon challenge. Detailed studies in this regard can help to increase the efficacy of the vaccine and might be implied to other vaccinations as well.

Towards this end, in the current study, we investigated transcriptomics profiles of pre-and postvaccination patients after the first, 2nd, and 3rd vaccination dose of RTS,S using various bioinformatics techniques, i.e., PCA, differential gene expression analysis, Hierarchical clustering, etc. Exploratory data analysis based on the PCA clearly shows distinct clusters of samples of control and CHMI. Subsequently, differential gene expression analysis using the edgeR scrutinized 219 significantly differentially expressed genes (FDR <0.005). Eventually, the biological role of significant genes was delineated using gene enrichment analysis.; which reveals the regulation status of chemokines and cytokines in pre-vaccinated and post-vaccinated samples at different doses. Gene enrichment analysis showed that *CCL8*, *CXCL10*, *CXCL11*, *XCR1*, *CSF3*, *IFNB1*, *IFNE*, *IL12B*, *IL22*, *IL6*, and *IL27* were involved in cytokine-chemokine pathways and upregulated after earlier doses but downregulated after the 3rd dose. Analysis of downregulated genes in DAVID and KEGG pathway illustrated how significantly downregulated genes after 3rd vaccination on CHMI is primarily associated with cytokine and inflammation. Over-expression of those inflammation and cytokine gene has been one of the driving forces of death and contribute to pathogenicity. [25] In fact, Chemoattractant cytokines or chemokines have proven to be regulators of leukocyte trafficking and potentially contribute to severe malaria. [37] [38] Our analysis suggested repeated doses of malaria vaccination help in protection because it is associated with the balanced expression of pro and anti-inflammatory cytokines. *SYT4*, *CBARP*, *NCS1*, *CACN1G*, *RHBDL3*, *CBARP*, etc., significant genes in our analysis have shown functions for calcium ion receptors, calcium gated ion channels, or calcium ion-dependent exocytosis according to gene ontology studies. [53–56] It has been observed that antibody levels to the voltage-gated calcium channels, but not to other ion channels, increase with the severity of malaria infection. [61]

Significantly elevated pro-inflammatory IL-6, IL-12 have been observed in the severe-malaria group compared to age-matched healthy children. [45] Also, IL12 has been found to induce IFN-γ, a key mediator of inflammatory immune responses. [42] IL-12 has shown evidence to play an essential role in the pathogenesis of malaria. [42] Furthermore, levels of IL-6 are elevated in malaria disease and contribute to disease severity. [31] [43] [44] Moreover, IFNB1-regulated genes were observed in severe cerebral malaria. [46] IL-12B was the most down-regulated gene after the 3rd dose in our study. Furthermore, TNF, IL-1, IL-6, etc., inflammatory genes are over-expressed in falciparum malaria. [30] [31]. These genes are associated with cardiac insufficiency and myocardial dysfunction. [32] [33] [34] Additionally, inflammatory/inducible chemokines CXCL10, CXCL11, and CCL8 suggest involvement in response to the malaria infection. [47]. Relative involvement of CXCL10 and CXCL11 has been found to recruit inflammatory leukocytes of malaria-infected mice. [37] Again, studies have indicated that the concentration of CXCL11 was greater in symptomatic than asymptomatic malaria and was upregulated among the fever-positive groups, which identified CXCL11 as a possible biomarker for malarial fever. [41] The activity and capacity of cytokines through directing sequestration and driving anemia contribute to restricting oxygen supply to mitochondria and making falciparum malaria primarily a cytokine-driven inflammatory disease. [41][25]

One of the pathways involved with the significant genes was, surprisingly, the cancer pathway. In Africa, malaria is known to influence genetic variation at several loci in the human genome [63], which might be involved in cancer and impact the biology and epidemiology of both diseases. [64] Moreover, recent evidence suggested that inflammatory cytokines might implicate several cancers. [65] Additionally, cancer–malaria interactions have been reported in the human liver where malaria parasites attack in its life cycle. [64] p53 protein has been observed to play a crucial role in hepatocyte infection by malaria parasite sporozoites. P53 is also the most highly mutated gene in several cancers. [66]. This area needs further investigation for better understanding as there is an excellent potential for new novel anti-cancer therapies using anti-malarial drugs.[67]

Further, many critical events in the Plasmodium life cycle are regulated by changes in the cellular levels of Ca2+ [51]. Moreover, levels of antibodies to the voltage-gated calcium channels correlate with the increased severity of malaria infection. [52] *SYT4*, *CBARP*, *NCS1*, *CACN1G*, *RHBDL3*, *CBARP*, etc., significant genes in our analysis have shown functions for calcium ion receptors, calcium gated ion channels, or calcium ion-dependent exocytosis according to literature study or gene ontology. [53–56] It has been observed that antibody levels to the voltage-gated calcium channels, but not to other ion channels, increase with the severity of malaria infection. [61] Moreover, P. falciparum protein PfRH1is found to trigger the release of calcium ions. Extensive involvement of calcium signaling has been observed in various crucial pathways of the parasite. Therefore, any interruption would be deleterious for invasion and, ultimately, the growth of the malaria parasite. So, components of calcium signaling are considered for therapeutic interventions. [62]

The malaria RTS,S/AS01 vaccine uses the CSP protein as a target antigen against malaria. [57] In earlier stages of vaccine development, higher concentrations of antibodies against CSP were observed on the day of the challenge. [59] Moreover, the control group without vaccination developed malaria earlier than the test group with three doses of vaccination, in which a strong humoral response against CSP was shown. [60] After the initial dose, there are low titers of antibodies against CSP protein. But after later doses, high antibody titers are present, and antibody feedback can further block immunodominant response. [67] Although inflammatory mediators have been repeatedly found to be implicated in the severity of the disease, this evidence gave rise to the widely held belief that severe malaria might be an immune-mediated disease. [5] [36] Also, parallels are present between sepsis and malaria associated with functionally crucial inflammatory cytokines present in both conditions. [25] Malaria-induced sepsis is related to an intense pro-inflammatory cytokinemia, though the mechanisms behind it are poorly understood. [35] Additionally, critical illness associating with an inflammatory response is a cause of multifactorial anemia. [28] Anemia could contribute to poor oxygenation of tissues in malaria patients. [29]

One of the adverse side effects of this is the over-expression of inflammatory genes, which often have the role of a double-edged sword. [58] As our study suggested, cell-mediated cytokines are less expressed after the 3^rd^ dose, which was not the case after the 1^st^ or 2^nd^ dose, confirming that multiple doses of malaria RTS,S vaccination is crucial in gene expression and control of inflammation in malaria infection. Downregulation of cytokine and inflammation-related genes helps decrease the negative side-effects of cytokine storms in malaria as excessive pro-inflammation increases the severity of malaria [25]. Therefore, cytokine overexpression as a part of humoral immunity was reduced after repeated vaccination doses, resulting in protection by reducing the complexities of malaria infection. Additionally, a fractional booster dose initiates high protection upon challenge by increasing antibody somatic hypermutation [10]. Moreover, exploratory data analysis showed evidence that protected-day 14 samples were distinct from other groups, which needs further investigation for early detection of malaria vaccination efficacy before the challenge. [supplementary figure 7]. Our study also suggests the necessity to further explore passive immunization with monoclonal antibodies as a new approach to prevent and eliminate malaria. [30] As cytokines are associated with various inflammatory diseases, the study of malaria vaccination’s control over inflammatory cytokine gene expression might become helpful in other diseases too, where they play a significant role in pathogenesis. [49]

## Conclusion

This study was able to identify 13 inflammatory genes whose expression in malaria vaccination played a significant role in the cytokine-cytokine receptor interaction pathway, JAK-STAT signaling pathway, and pathways in cancer. Furthermore, we demonstrated a comparatively less focused protection mechanism after vaccination and discussed the gene expression pattern of various vaccination doses. We analyzed the dual role of protection and pathogenicity of cytokines in malaria infection and how multiple doses of vaccination increase protection by influencing these cytokine levels and producing antibodies against the malaria CSP antigen.

## Supporting information

Supplementary Figure 1:Differential gene expression analysis using edgeR Pipeline

Supplementary Figure 2: Factor Regression Pipeline

Supplementary Figure 3: PCA plot for control vs. CHMI samples for significant genes

Supplementary Figure 4: Significantly expressed genes in males and females on control vs CHMI samples

Supplementary Figure 5: JAK-STAT pathway

Supplementary Figure 6: Pathways in cance

Supplementary Figure 7: PCA plot of Protected and Non-Protected samples on various days

Supplementary Table 1: Significantly expressed genes in male and female on control vs CHMI samples.

Supplementary Table 2: Significantly expressed genes in protected and non-protected groups on control vs. CHMI samples.

## Authors Contribution

Supantha Dey performed the data analysis and produced the figures and wrote the manuscript. Harpreet Kaur reviewed the data, provided technical assistance with analysis, drafted the manuscript, and supervised the whole project. Elia Brodsky and Dr. Mohit Mazumder provided expert reviews of the project.

## ACKNOWLEDGMENTS

I thankfully acknowledge Pine Biotech for helping me with their efforts on this project. The groundwork of this project was completed during the Omics Logic Fellowship Program. The processing and visualization pipelines were generated with the T-BioInfo Server.

## Abbreviation

RBC: Red Blood Cell
CHMI: Controlled Human Malaria Infection
Ch: Day of Challenge
P.V.: Pre-vaccination
PfRH1: Plasmodium falciparum reticulocyte binding protein homolog 1
ChF/ChM: Female or Male samples on CHMI day
PreF/PreM: Female or Male samples on pre-challenge or control samples
PCA: Principal Component Analysis

