## Supplementary figures and images for "Analysis of Gene Expression Profiles to study Malaria Vaccine Dose Efficacy & Immune Response Modulation"

### Supplementary Figure 1:Differential gene expression analysis using edgeR Pipeline

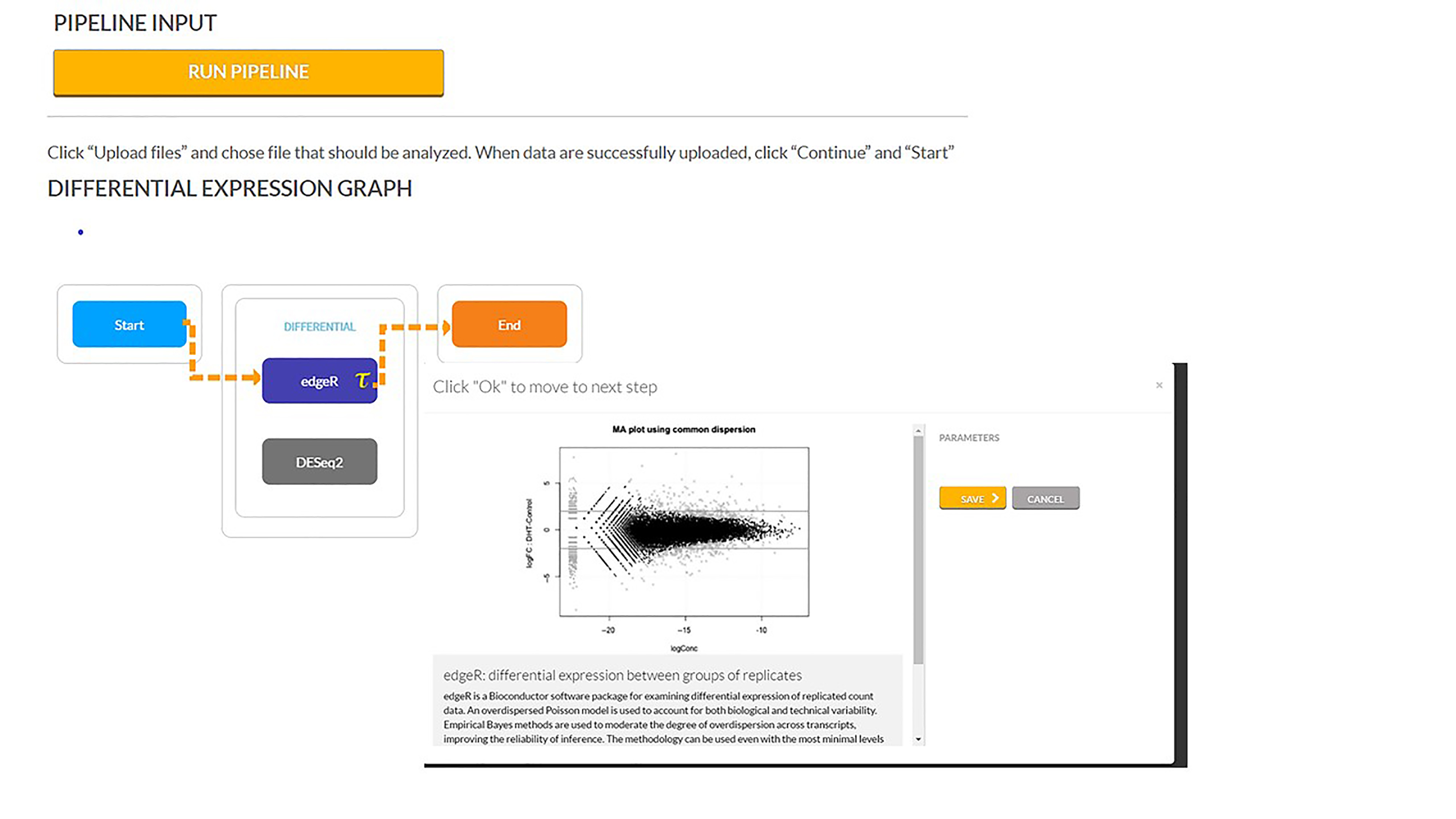

### Supplementary Figure 2: Factor Regression Pipeline

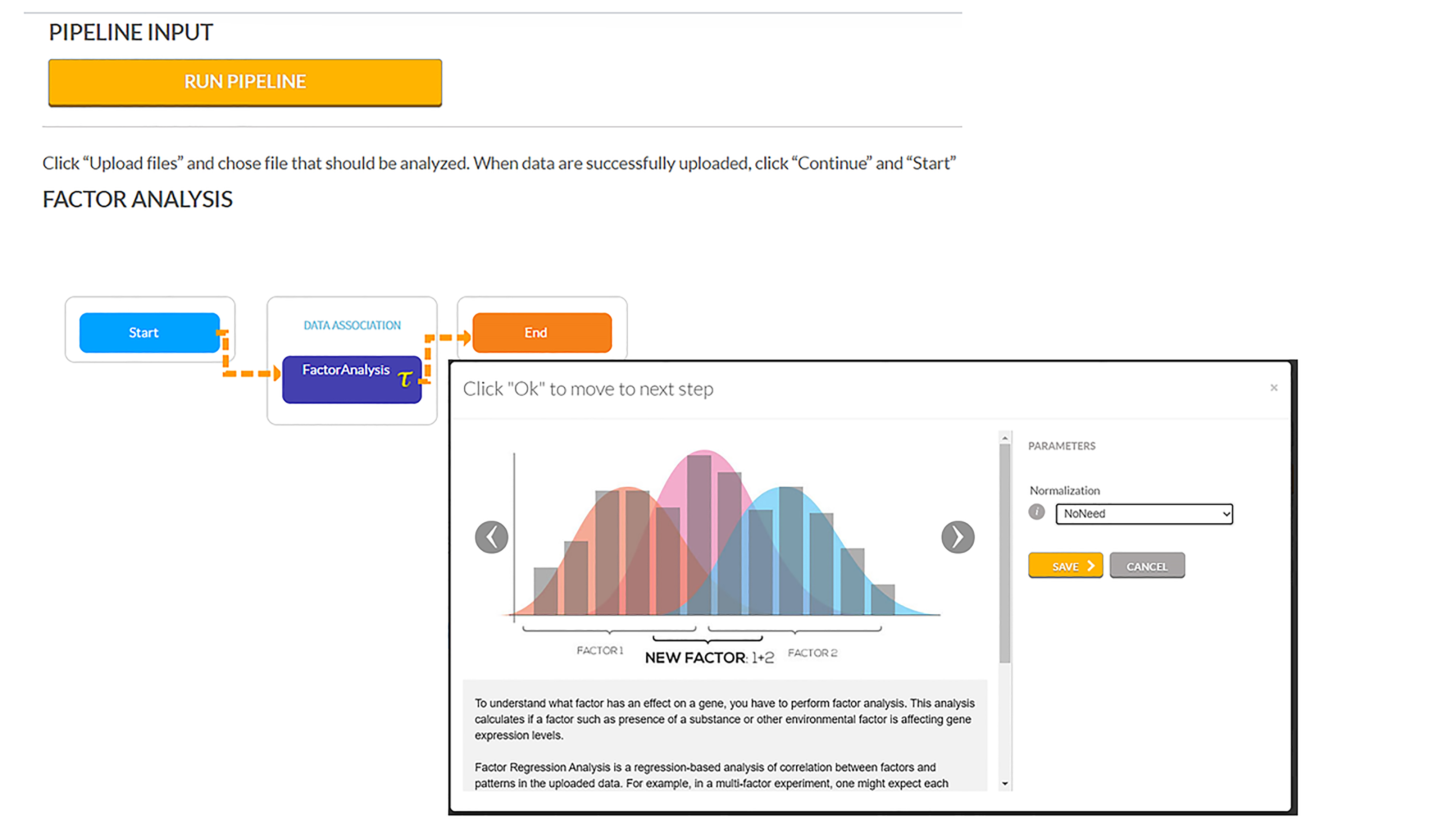

### Supplementary Figure 3: PCA plot for control vs. CHMI samples for significant genes

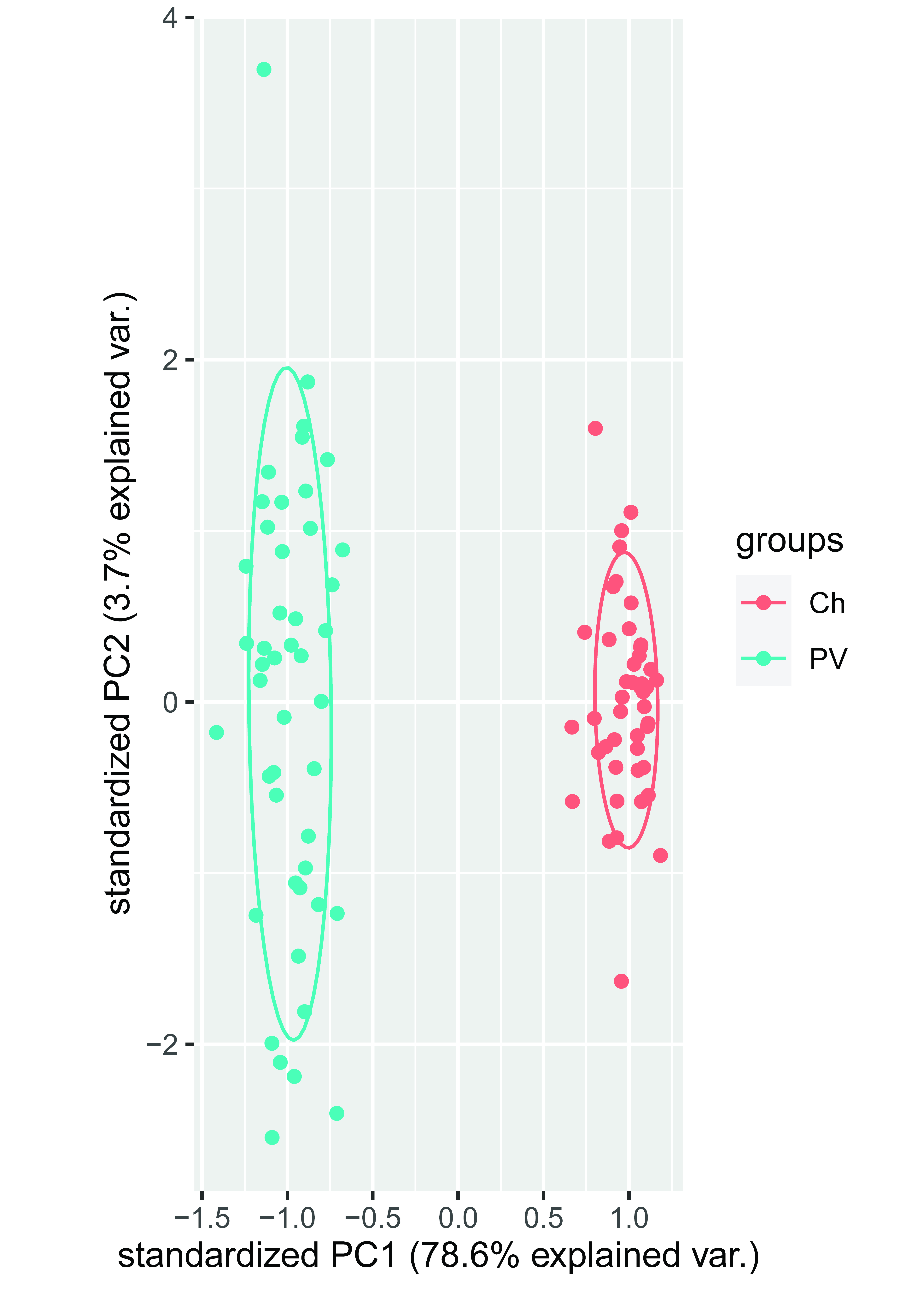

### Supplementary Figure 4: Significantly expressed genes in males and females on control vs CHMI samples

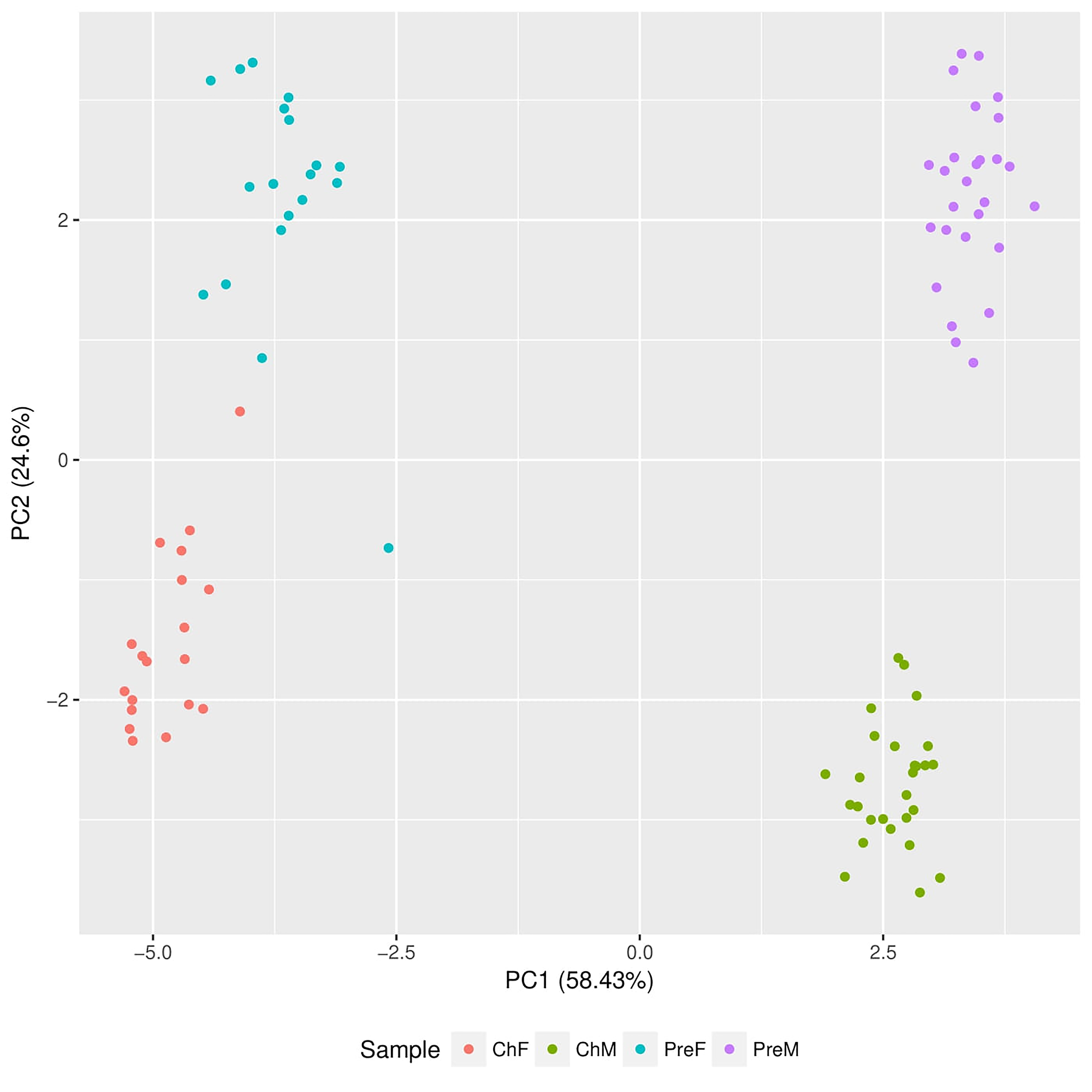

### Supplementary Figure 5: JAK-STAT pathway

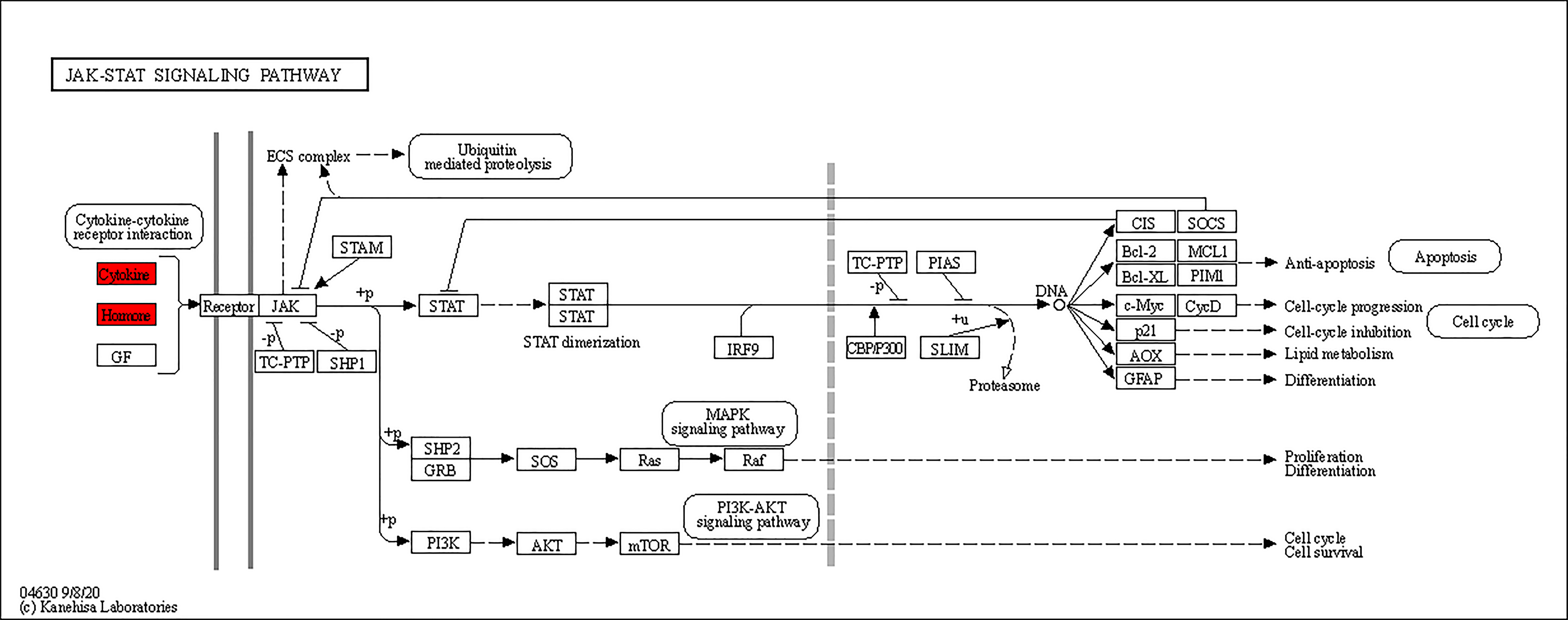

### Supplementary Figure 6: Pathways in cance

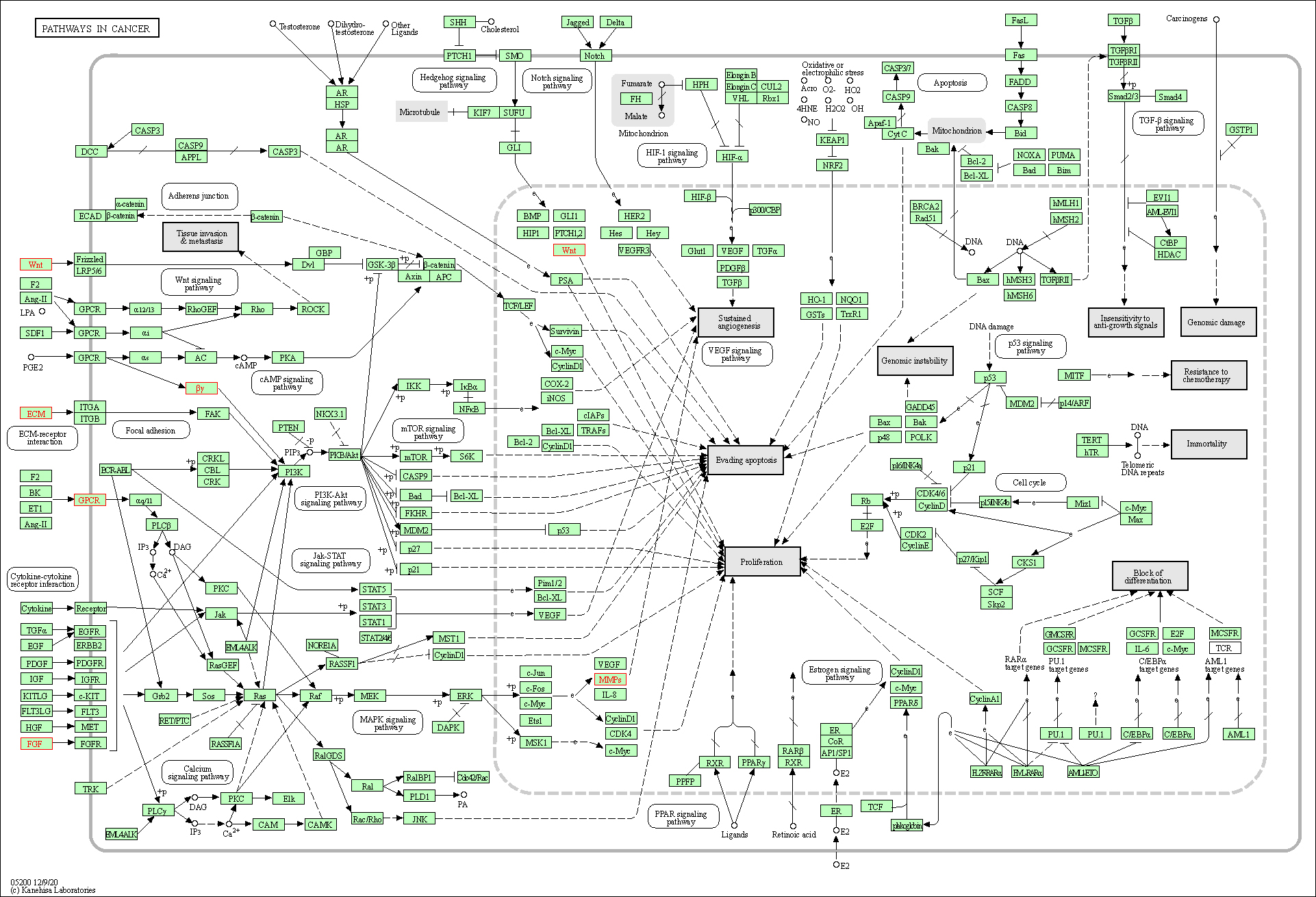

### Supplementary Figure 7: PCA plot of Protected and Non-Protected samples on various days

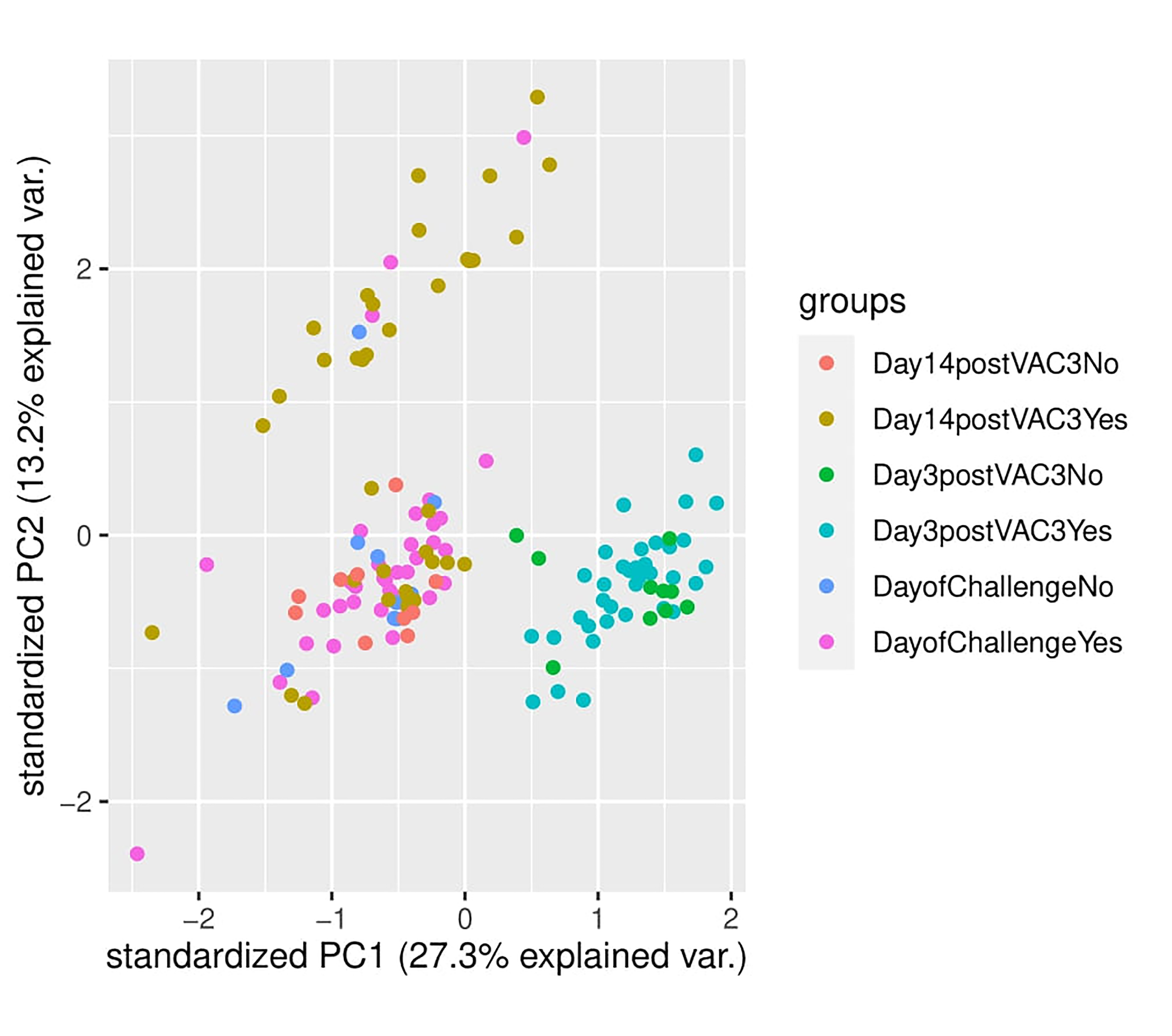
