## Supplementary Table 1: Significantly expressed genes in male and female on control vs CHMI samples. for "Analysis of Gene Expression Profiles to study Malaria Vaccine Dose Efficacy & Immune Response Modulation"

| Ensembl ID | GeneSymbol |
| --- | --- |
| ENSG00000108700 | CCL8 |
| ENSG00000148053 | NTRK2 |
| ENSG00000151012 | SLC7A11 |
| ENSG00000160097 | FNDC5 |
| ENSG00000163879 | DNALI1 |
| ENSG00000184368 | MAP7D2 |
| ENSG00000197980 | LEKR1 |
| ENSG00000231535 | LINC00278 |
| ENSG00000233864 | TTTY15 |
| ENSG00000240184 | PCDHGC3 |
