## Supplementary Table 2: Significantly expressed genes in protected and non-protected groups on control vs. CHMI samples. for "Analysis of Gene Expression Profiles to study Malaria Vaccine Dose Efficacy & Immune Response Modulation"

| Ensembl ID | GeneSymbol |
| --- | --- |
| ENSG00000074410 | CA12 |
| ENSG00000100065 | CARD10 |
| ENSG00000136944 | LMX1B |
| ENSG00000145506 | NKD2 |
| ENSG00000151952 | TMEM132D |
| ENSG00000162745 | OLFML2B |
| ENSG00000163666 | HESX1 |
| ENSG00000169507 | SLC38A11 |
| ENSG00000182901 | RGS7 |
| ENSG00000197993 | KEL |
| ENSG00000213934 | HBG1 |
| ENSG00000222414 | RNU2-59P |
| ENSG00000224730 | LILRB1-AS1 |
